# Corticospinal correlates of hand preference for reaching during whole-body motion

**DOI:** 10.1101/2021.01.05.425205

**Authors:** Leonie Oostwoud Wijdenes, Syanah C. Wynn, Béla S. Roesink, Dennis J.L.G. Schutter, Luc P.J. Selen, W. Pieter Medendorp

## Abstract

Behavioral studies have shown that humans account for inertial acceleration in their decisions of hand choice when reaching during body motion. Physiologically, it is unclear at what stage of movement preparation information about body motion is integrated in the process of hand selection. Here, we addressed this question by applying transcranial magnetic stimulation over motor cortex (M1) of human participants who performed a preferential reach task while they were sinusoidally translated on a linear motion platform. If M1 only represents a read-out of the final hand choice, we expect the body motion not to affect the MEP amplitude. If body motion biases the hand selection process prior to target onset, we expect corticospinal excitability to modulate with the phase of the motion, with larger MEP amplitudes for phases that show a bias to using the right hand. Behavioral results replicate our earlier findings of a sinusoidal modulation of hand choice bias with motion phase. MEP amplitudes also show a sinusoidal modulation with motion phase, suggesting that body motion influences corticospinal excitability which may ultimately reflect changes of hand preference. The modulation being present prior to target onset suggests that competition between hands is represented throughout the corticospinal tract. Its phase relationship with the motion profile suggests that other processes after target onset take up time until the hand selection process has been completely resolved, and the reach is initiated. We conclude that the corticospinal correlates of hand preference are modulated by body motion.

---

We frequently encounter tasks that can be performed with either hand, for example moving papers on a desk, picking up a key from the table, or opening a door. Whether we use our left or right hand is known to depend on various factors, including handedness, recent choice success, and eye and head position (Bakker et al. 2018; Schweighofer et al. 2015; Stoloff et al. 2011). Biomechanical factors also play a role: participants prefer to move the hand that is closest to the target (Mamolo et al. 2004; Przybyla et al. 2013) and for two equidistant targets participants choose to move to the target that can be reached with the lowest biomechanical cost (Cos et al. 2011, 2014).

Recently, Bakker et al. (2017, 2019) studied hand choice when participants are in motion. In such a dynamic situation, not only vision and proprioception provide information about the state of the body and the environment, also information about whole body motion is registered by the vestibular organ (Angelaki and Cullen 2008). Full-body acceleration differentially modulates the biomechanical costs of left and right-hand movements and consequently hand preferences are modulated by the current dynamic situation (Bakker et al. 2017, 2019). The physiological basis of this motion related modulation of hand preference is unknown.

It has been proposed that decision making and movement generation processes are tightly connected in the sensorimotor areas of the brain (Cisek 2007; Cisek and Kalaska 2010). For hand selection, this implies that motor plan for both hands are generated in parallel, while these two plans compete for execution. It is unclear at what level this competition between the two motor plans is resolved.

On the one hand, studies suggest that competition for hand selection is resolved before movement preparation reaches dorsal premotor cortex (PMd), possibly in parietal cortex (Bernier et al. 2012; Dekleva et al. 2018; Oliveira et al. 2010). On the other hand, it has been observed that areas closer to movement execution, up to primary motor cortex (M1), represent evidence for multiple concurrent movements (Cisek and Kalaska 2005; Dekleva et al. 2016; Derosiere et al. 2019; Thura and Cisek 2014).

Transcranial magnetic stimulation (TMS) over the motor cortex can be used to obtain a non-invasive physiological read-out of the state of corticospinal excitability, as evaluated by electromyographic recordings of the motor-evoked potential (MEP) (Bestmann and Krakauer 2015). In preferential reaching tasks, corticospinal excitability is enhanced for the selected hand, while it is suppressed for the non-selected hand (Duque et al. 2010; Duque and Ivry 2009; Klein-Flügge et al. 2013; Klein-Flügge and Bestmann 2012). Here we examine if M1 represents the modulation of hand preference with full-body motion by applying a single TMS pulse over M1 to quantify corticospinal excitability at the moment a reach target would have been presented. In this way, we learn how full-body motion affects hand preference.

We hypothesized that if M1 only represents a read-out of an already made decision for which the competition was resolved in upstream areas, corticospinal excitability would not be modulated by the whole-body motion if no target is presented. However, if body motion affects hand preference prior to target onset, we expected corticospinal excitability to modulate dependent on the whole-body motion, even before a target is presented. Corticospinal excitability was indexed by the MEP amplitude of the contralateral lateral triceps muscle.

## Methods

### Participants

20 self-reported right-handed participants (15 females) aged 19-47 years old (mean age 25 years) took part in this study, consisting of an intake session and two experimental sessions. Participants had normal or corrected-to-normal visual acuity, and had no history or presence of neurological or psychiatric disorders by self-report. Due to technical problems, data of one female participant had to be discarded. Participants received written and verbal information about the study prior to providing written informed consent, whereby they remained naïve as to the research question. Participants refrained from taking psychotropic substances within two hours prior to experimentation and from taking alcohol within 24 hours prior to experimentation. This study was approved by the medical research ethics committee of the Radboud University Medical Center Nijmegen (NL59818.091.16).

### Apparatus

Participants were seated on a vestibular sled in a darkened room (Figure 1A). The sled was powered by a linear motor (TB15N; Technotion, Almelo, The Netherlands) and controlled by a Kollmorgen S700 drive (Danaher, Washington, DC, USA). Participants were securely fastened with a five-point seat belt. Their head was immobilized with a personalized thermoplastic mask (Posicast). Visual stimuli were presented on a 27-inch touch screen which also registered touch of the two index fingers (ProLite; Iiyama, Tokyo, Japan). The position of both index finger tips and the sled were measured at 500 Hz using an Optotrak Certus system (Northern Digital, Waterloo, Canada). Electromyographic activity of six right arm muscles was recorded using a Trigno Wireless EMG system (Delsys, Boston, USA): first dorsal interosseous, brachioradialis, biceps long head, biceps short head, triceps lateral head (TLAT) and triceps long head. EMG data were band-pass filtered (30-450 Hz), amplified (1000) and sampled at 1111 Hz.

**Fig 1.**
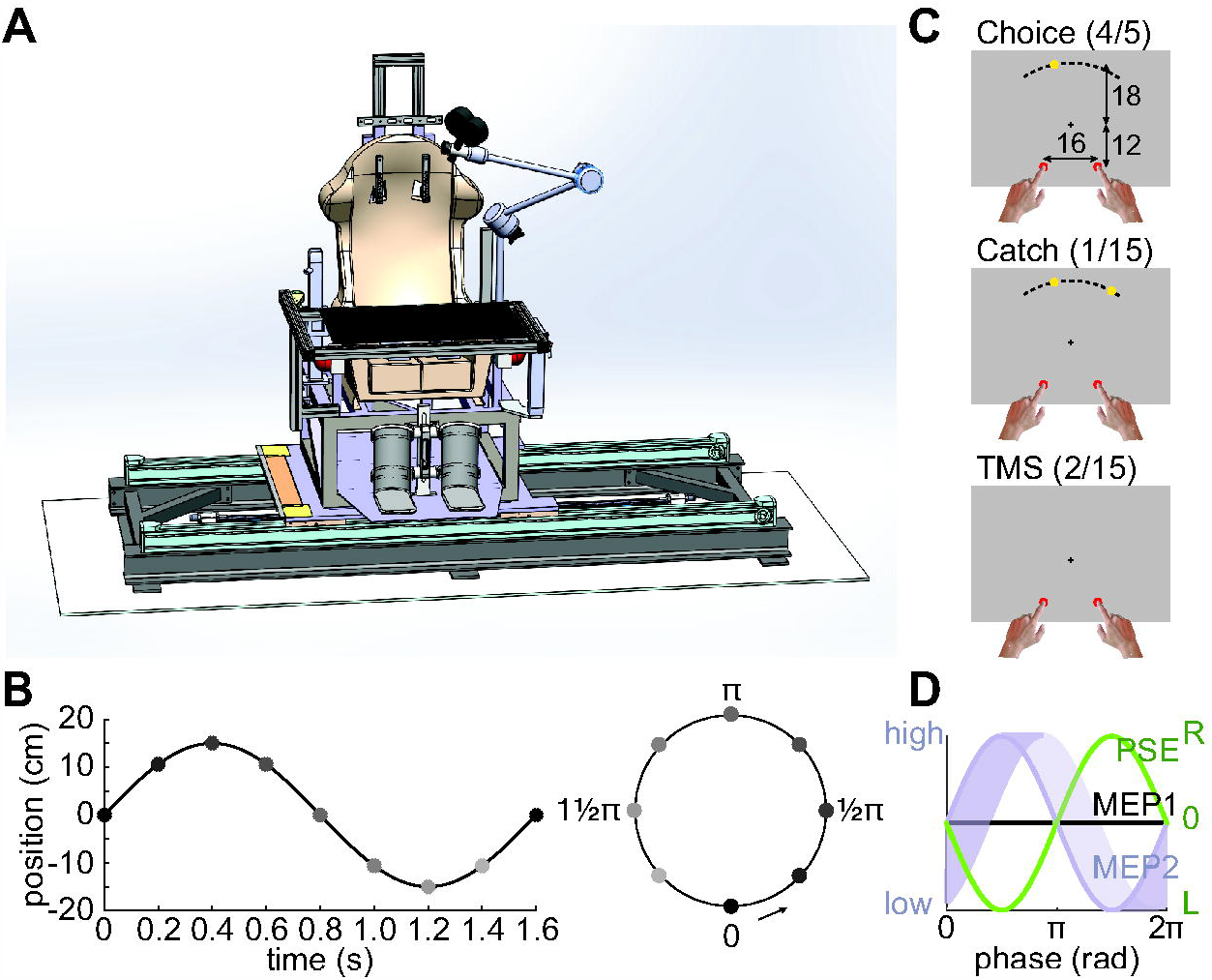
Experimental set-up. **A**. Illustration of the vestibular sled, touch screen and TMS coil. **B**. Sled position as a function of time. Target stimuli were presented, or TMS stimulation was applied at one of eight phases of whole-body motion (grey circles). **C**. Start locations of the index fingers (red circles), fixation cross and example target locations (yellow circles) for choice, catch and TMS trials. **D**. Predictions for the modulation of hand preference and corticospinal excitability as a function of sled phase. Based on Bakker et al. (2017) we expect maximum leftward deviation of the PSE at maximum leftward acceleration, i.e. at phase ½ pi (green). MEP may (MEP2, purple) or may not (MEP1; black) modulate as a function of sled phase. The shaded area for MEP2 indicates the predicted corticospinal excitability for a read-out in the time window from target presentation to movement initiation.

For the MEP measurements we targeted the TLAT muscle, as this is the primary actor of the reaching movement. To elicit MEPs, a figure-of-8 coil (Cool-B65, MagVenture A/S) was placed over left M1 to target the right arm TLAT. The coil was oriented posterolaterally at an angle of ∼45° to the midline and fixed to the sled. The coil was securely fastened to the sled. Together with the mask this configuration ensured that there was minimal motion between the coil and the head within a session. Stimulation parameters were in agreement with the International Federation of Clinical Neurophysiology safety guidelines (Rossi et al. 2009). There were no serious adverse events and participants had no issues tolerating the TMS. Since TLAT is the primary actor of the reaching movement and this was the targeted muscle, only data from TLAT will be reported.

### Experiment

The intake session and two experimental sessions took place at different days and all started with localizing the right arm TLAT hotspot and determining the resting motor threshold for this muscle (Schutter and van Honk 2006). If we could not elicit a MEP at a stimulation intensity 83% (as a percentage of the maximum machine output), or if the participant did not feel comfortable with the experimental setup, volunteers were not invited to take part in the experimental sessions. Therefore, we saw about three times as many volunteers in the intake session than volunteers who took part in the full experiment. The mean resting motor thresholds in the experimental sessions of the participants who completed the experiment was 70.2% (*SD* = 11.3) of the maximum machine output. These relatively high motor thresholds are probably related to the targeted muscle. After the resting motor threshold was determined in the intake session, participants were familiarized with the experimental setup and fitted with the personalized head mask.

During the experimental sessions, the sled translated in a sinusoidal fashion along the interaural axis with an amplitude of 0.15 m and a period of 1.6 s (Figure 1B), resulting in a peak velocity of 0.59 m/s and peak acceleration of 2.3 m/s^2^. While in motion, participants looked at a fixation cross and triggered the start of each trial by placing their left and right index fingers on the starting points (red circles, 3.5 cm diameter; Figure 1C). There were three types of trials: choice trials, catch trials and TMS trials (Figure 1C). In choice trials, a target was presented (yellow circle, 3.5 cm diameter) at one of eight phases of the whole-body motion (grey circles in Figure 1B).

Targets appeared within 5° of the intended phase of sled motion. In 75% of the trials, the direction of the presented target was determined by a Bayesian adaptive approach in order to find the target angle for which participants were equally likely to choose their left and right hand (Kontsevich and Tyler 1999; Prins 2013), whereby possible angles were −40°, −35°, −30° to 30° with steps of 2°, 35° and 40°. In the other 25% of trials, a peripheral target (−40°, −35°, −30°:2:-22°, 22°:2:30°, 35°, or 40°) was presented, enabling an estimate of the full psychometric curve after data collection. The adaptive estimation was run for each phase of motion separately. Participants were instructed to hit the target as quickly and accurately as possible with either their left or their right index finger. In catch trials, to avoid pre-determined hand choices, two targets were presented and participants were instructed to hit both targets with their left and right index fingers.

In TMS trials, a single TMS pulse (∼ 1 ms) at 120% of the participants’ resting motor threshold was delivered at one of eight phases of motion (grey circles in Figure 1B). Thus, the pulse was delivered at the time a target would have been presented in a choice trial, but the target remained absent in the TMS trials. After a TMS trial, there was a 3 s break and participants were asked to lift their fingers and replace them at the start locations. Trial type was pseudo-randomized whereby there were at least three other trials in between successive TMS or catch trials. Per session participants performed 6 blocks of 120 trials with short breaks in between the blocks. Each block consisted of 96 choice, 16 TMS and 8 catch trials, resulting in a total of 1440 trials per participant. Per phase of motion there were 24 TMS trials. One experimental session tested at the phases of sled motion 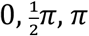 and 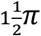, and the other session tested at 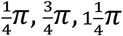 and 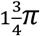. The order was counterbalanced across participants.

### Analyses

Hand choice was determined by the first index finger leaving the touch screen, as registered online by the screen. Optotrak data confirmed the choices determined based on touch screen data. For each sled phase, the target angle for which participants were equally likely to choose their left and right hand was estimated by a cumulative Gaussian distribution fit using a maximum likelihood approach with a lapse rate (Wichmann and Hill 2001):

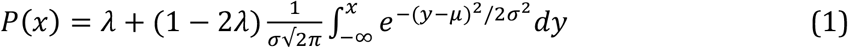

Here, *x* represents the target angle, *μ* represents the target angle for which participants were equally likely to choose their left and right hand, i.e. the point of subjective equality (PSE), *σ* represents the standard deviation of the choice distribution and *λ* represents the lapse rate.

Based on Bakker et al. (2017), PSE was expected to modulate with phase (Figure 1D; green). To determine the phase modulation of the sled on the PSE, two sinusoids with a coupled phase (*θ*_*PSE*_) and two independent amplitudes (A1 and A2) and offsets (B1 and B2) were fit to each participants PSEs of the two sessions:

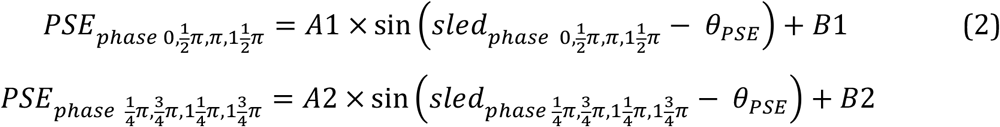

This ensured that differences in amplitude and offset, that may occur due to two different testing days, were accounted for.

Corticospinal excitability was determined by measuring the MEP amplitude caused by the single pulse TMS. For each trial, the difference between the maximum and minimum EMG activity in TLAT 15-35 ms after the TMS pulse was calculated (Cos et al. 2014). Trials were excluded if the maximum EMG activity in a window 200 ms before the TMS pulse exceeded 0.1 mV (Klein-Flügge and Bestmann 2012), if the trigger was missing or if sensor connection was lost. The trigger happened to be missing in one full session of participant 11. Of all other trials of all participants 9% was excluded. MEP was determined as the mean potential per participant per phase.

Similar to the hand choice data, to account for differences between the sessions in absolute stimulation intensity, MEP amplitude and offset, two sinusoids with a coupled phase (*θ*_*MEP*_) and two independent amplitudes (C1 and C2) and offsets (D1 and D2) were fit to each participants MEPs of the two sessions:

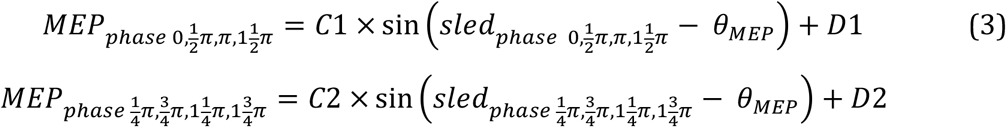

Since MEP is a noisy measure, the sinusoid fits were also performed on all single trial MEPs per participant instead of on the mean MEP per phase per participant. Also, a single sinusoid phase was fit to all participants mean MEPs with session and participant dependent amplitudes and offsets. All of these fits resulted in a similar estimation of the mean phase, suggesting that the measure is robust. Therefore, we only report results of the individual fits to the mean MEPs.

To test if there was a sinusoidal modulation of the PSEs and MEPs, or if a constant offset per session could better explain the behavioral and physiological data (see Figure 1D), a constant model was also fit to the data of each participant:

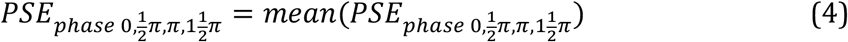

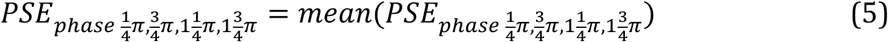

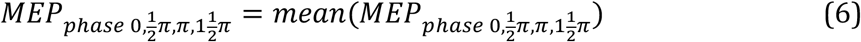

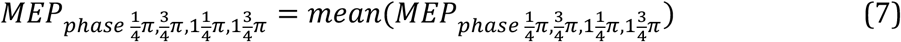

For every participant, the fits of the two models were compared by computing the Bayesian Information Criterion (BIC), which accounts for the difference in the number of parameters:

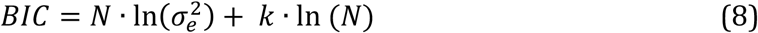

where *N* is the number of fitted data points, 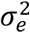 is the mean squared error of the fit and *k* is the number of model parameters, i.e. 1 for the constant model and 3 for the sinusoid model. The BIC value is smaller if the model has fewer parameters and hence provides a more parsimonious description of the data. To compare the two models a difference value was computed:

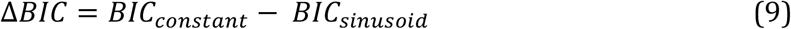

A BIC value difference of 2-6 indicates positive evidence for the model with the lower value, 6-10 indicates strong evidence and >10 very strong evidence (Kass and Raftery 1995).

Since MEPs were induced by stimulating left M1, we expected that MEPs would be enhanced for phases where a right hand choice was more likely. Behaviorally, a more likely right hand choice corresponds to a PSE shift towards the left (Figure 1D, green). If the modulation of MEP is aligned with the presentation of the target, we therefore expect a π phase difference between PSE and MEP (Figure 1D, purple). However, the modulation of MEP may not be aligned with target presentation, because information about the target may take some time to process in the brain, i.e. nondecision time (Ratcliff and McKoon 2008). Maximally, this process would last as long as the reaction time, which is ∼300 ms in this task (Bakker et al. 2017). With a sled period of 1.6 s, this would result in a 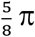 phase difference between PSE and MEP (Figure 1D, shaded purple). Thus, we hypothesize that the phase difference between PSE and MEP will be in between 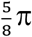 to π (Figure 1D). To test if there was a phase difference between PSE and MEP, we performed a Watson-Williams test (Berens 2009). To examine if the phase difference between PSE and MEP is in a congruent direction across participants we performed a correlation with a correction for circular data (Berens 2009).

## Results

We investigated if corticospinal excitability before a target is presented reflects biases in hand preference induced by whole-body motion. In most trials, participants were free to choose with which hand they preferred to move to the target. Figure 2A shows hand choice behavior of participant 9, separately for the different sled phases. Cumulative Gaussian fits were used to estimate the target angle for which participants were equally likely to choose their left and right hand, i.e. the PSE, indicated by the vertical black line. Figure 2B shows the PSE as a function of the sled’s motion phase at which the target was presented for the individual participants. Data from the two sessions are indicated by dark and light blue. To determine the phase relationship between sled motion and hand preference, the PSEs of each participant were fitted by two sinusoids with a single phase and session-dependent amplitudes and offsets (eq. 3). Consistent with previous work from our lab, the PSE was shifted mostly to the left, and thus indicating a preference for using the right hand, around maximum leftward acceleration, i.e. sled phase 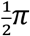. Similarly, the PSE was shifted most strongly to the right around maximum rightward acceleration, i.e. sled phase 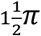, (Bakker et al. 2017, 2019).

**Fig 2.**
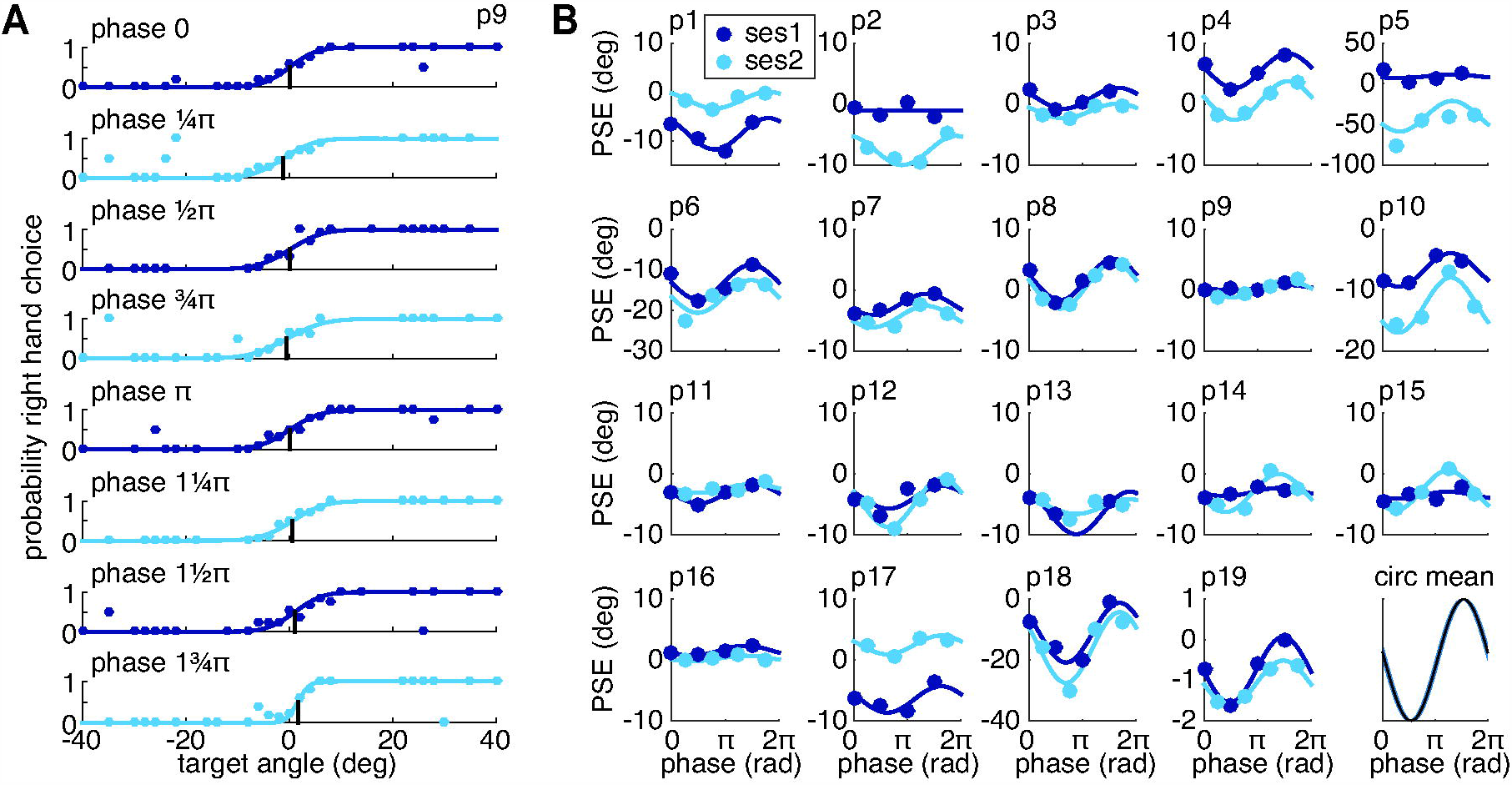
Choice behavior. **A**. Probability of right hand choice as a function of target angle (dots) fit by a cumulative Gaussian distribution (lines) for participant 9. The vertical black line indicates the PSE angle. Each panel shows a different sled phase. **B**. PSE as a function of sled phase for all participants (p1 to p19) and the circular mean phase with SEM (bottom right panel). PSEs were tested in two different sessions (dark blue: 0, ½π, π, 1½π; light blue: ¼π, ¾π, 1¼π, 1¾π). Lines show sinusoidal fits with a within participant coupled phase and session dependent amplitudes and offsets (eq. 2).

To test if the PSE data is better represented by a sinusoid than by a constant offset, we calculated the difference in BIC between the two models, thereby accounting for the difference in number of free parameters. As illustrated in Figure 3, left panel, 16 out of 19 participants show strong to very strong evidence (Δ*BIC* > 6) for a sinusoidal modulation, while no participant shows positive evidence for a constant offset (Δ*BIC* < − 2). This again confirms that hand choice is modulated by sinusoidal body motion in a sinusoidal fashion. The modulation is thought to reflect the influence of bottom-up acceleration signals on hand choice (Bakker et al. 2017, 2019).

**Fig 3.**
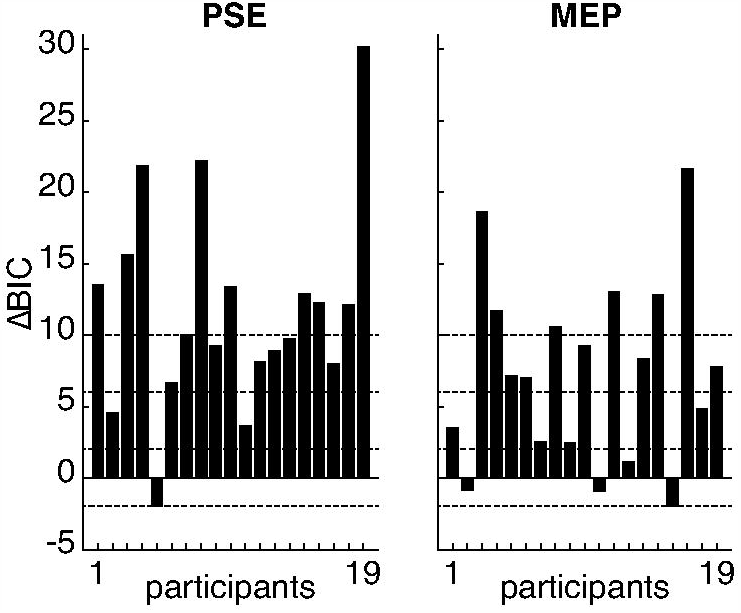
BIC model comparisons for the PSE (left panel) and MEP data (right panel). Δ*BIC* values per participant. A positive value indicates support for the sinusoidal model over the constant model.

In selected trials a single TMS pulse was delivered. Figure 4A shows the mean TLAT EMG response (MEP) to this pulse for each sled phase for participant 9. Figure 4B shows the resulting MEP amplitudes as a function of sled phase for all participants. Compared to the PSEs, MEPs were more variable between sessions and between participants. As for the PSEs, a sinusoidal model with a single phase and session dependent amplitudes and offsets was fit to the MEPs for each participant (eq. 3). Across participants, the circular mean phase seems to peak around phase π (bottom right panel).

**Fig 4.**
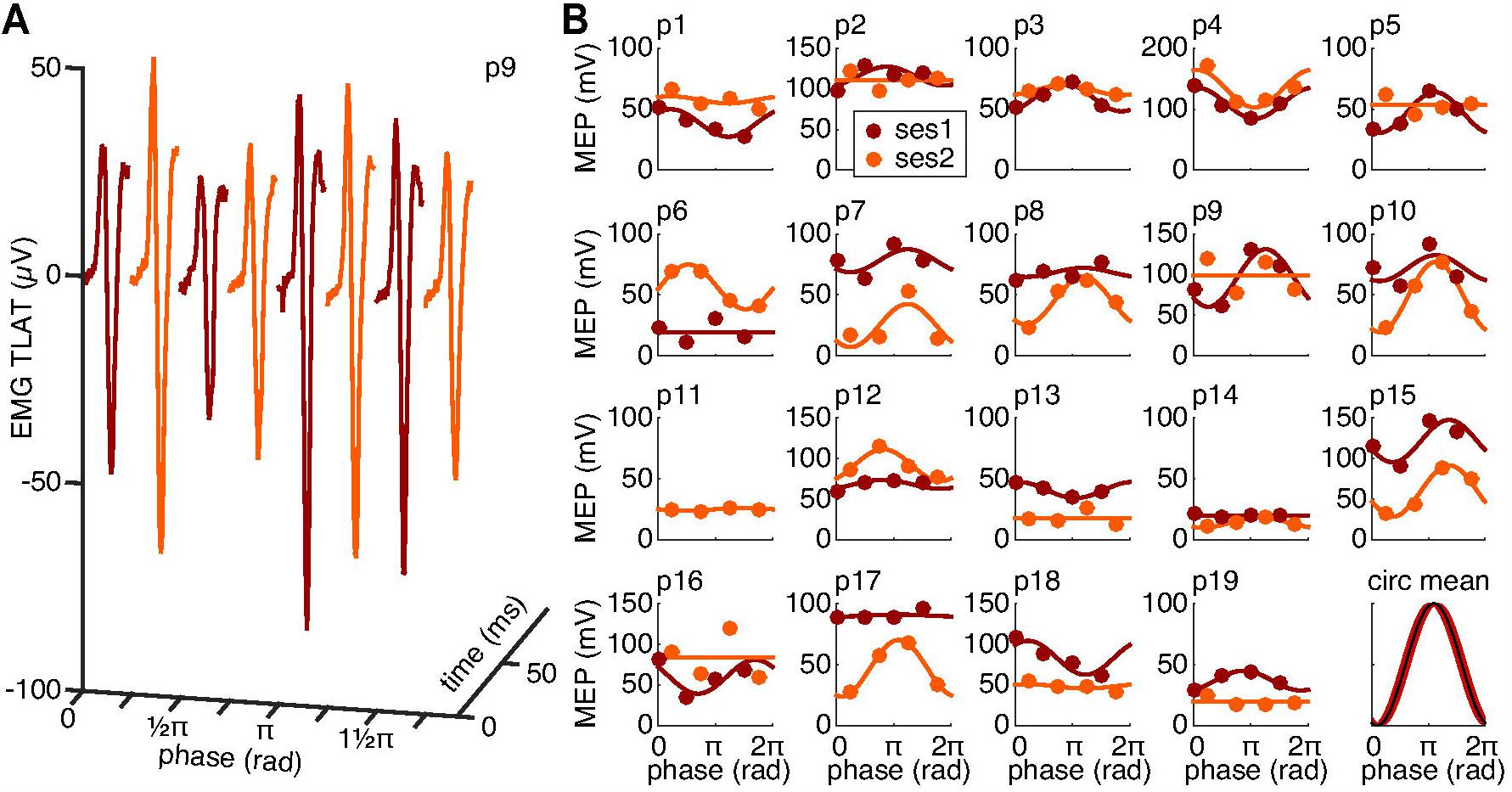
Corticospinal excitability. **A**. Time as a function of average EMG response of TLAT for participant 9. Each panel shows a different sled phase. **B**. Mean MEP amplitude as a function of sled phase for all participants (panels 1:19) and the circular mean phase with SEM (bottom right panel). MEPs for each phase were tested in two different sessions (dark red: 0, ½π, π, 1½π; orange: ¼π, ¾π, 1¼π, 1¾π). Lines show the sinusoid fits with a within participant coupled phase and session dependent amplitudes and offsets.

To test if the MEP data, similar to the PSE data, also support a sinusoidal model over a constant offset, the difference in BIC value between the two models was calculated (Figure 3, right panel). Here, 15 out of 19 participants show positive to very strong evidence (Δ*BIC* > 2) for a sinusoidal modulation, while no participant showed positive evidence for a constant offset (Δ*BIC* < −2). This suggests that the MEPs were modulated by the full body motion in a sinusoidal fashion.

We hypothesized that if corticospinal excitability reflects biases in hand preference, there would be a 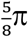 to π phase difference between the PSE and MEP phase (Figure 1D). Figure 5A shows a polar plot of the PSE and MEP phases for each participant. A Watson-Williams test indicated that there was a significant phase difference between PSE (M = 3.3 rad) and MEP (M = 1.9 rad) (*F*(1,36) = 11.97; *P* = 0.0014). The difference between the mean phases is 1.4 rad.

**Fig 5.**
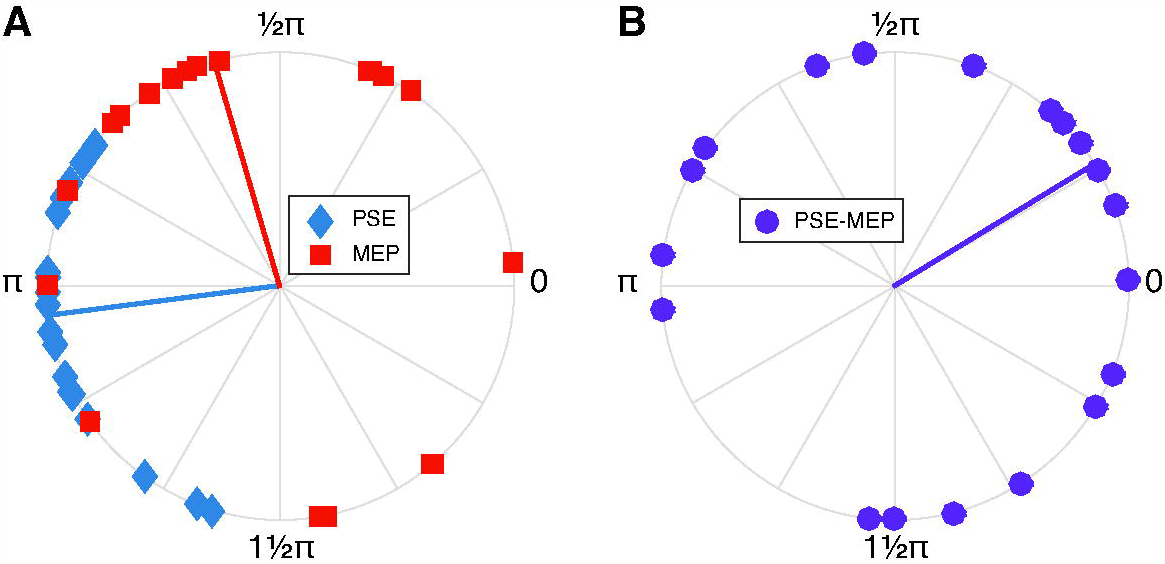
Correlation between PSE and MEP. **A**. Estimated PSE (blue diamonds) and MEP (red squares) phase in radians for all participants. Lines indicate the circular mean phase. **B**. Circular phase difference between PSE and MEP for all participants (circles) and the across participants mean phase difference (line) in radians.

Figure 5B shows the phase difference between PSE and MEP phase for each participant. Note that there is a discrepancy between the difference of the means (lines in Figure 5A) and the mean of the individual differences (line in Figure 5B) due to the circular nature of the data. A circular correlation indicated that the difference between PSE and MEP phase is consistent across participants (*R* = −0.5, *P* = 0.0141), with on average PSE phase leading MEP phase by 0.55 rad. Although both phase differences are in the expected direction, the difference seems to be just outside the hypothesized window.

## Discussion

We examined if corticospinal excitability reflects hand choice preference due to whole-body motion before the hand is selected. Choice behaviour confirms previous observations from our lab: sinusoidal whole-body motion modulates hand choice bias. Specifically, the target for which both hands are equally likely to be selected shifts maximally to the left (indicating a preference for using the right hand) at maximum leftward body acceleration (Figure 2B), and maximally to the right at maximum rightward body acceleration (Bakker et al. 2017, 2019). Corticospinal excitability also modulates sinusoidally with body motion. Stimulation over left M1 resulted in maximum excitability around phase π (maximum leftward body velocity, Figure 4B). The sinusoidal modulation of corticospinal excitability suggests that biased competition between hands penetrates deeply within the motor system. This fits within a framework of multiple concurrently prepared actions, even before a target is presented (Cisek and Kalaska 2005; Derosiere et al. 2019; Thura and Cisek 2014).

The fact that both the hand choice bias and MEP amplitude are sinusoidally modulated by whole body motion, raises the question whether the MEP modulation is predictive to the hand preference. MEPs were elicited at the moment that otherwise a target would be presented. If hand preference is reflected in the corticospinal state at the moment of target presentation, we would have expected a phase difference between PSE and MEP of *π*: a maximum shift of the PSE to the left (negative) corresponds to a maximum MEP amplitude (Figure 1D). However, if behavioral choice is influenced by the corticospinal state somewhere in the reaction time window, the phase of the MEP modulation would shift further along the sled motion (to the right in Figure 1D; sled motion period is 1.6 s), resulting in a smaller phase difference between PSE and MEP. This MEP phase shift would maximally last as long as reaction time (∼300 ms), resulting in a phase difference between PSE and MEP of 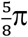. The mean phase difference that we found was even slightly smaller than the hypothesized window. This might suggest that hand preference is not fully predictable by corticospinal excitability before a target is presented.

It has been shown that vestibular information is taken into account in movement planning and online control of reaching movements. Vestibular stimulation by means of body rotation, or by means of artificial stimulation of the vestibular organ with galvanic stimulation which induces the illusion of a body rotation, results in corrections of the reaching movement to account for the perceived rotation (Bresciani et al. 2005; Keyser et al. 2017; Oostwoud Wijdenes et al. 2019; Reichenbach et al. 2016). Also, visuomotor feedback gains for online corrections are modulated by vestibular information (Oostwoud Wijdenes et al. 2019).

In hand choice tasks with a stationary body, biomechanical costs in terms of required effort have been shown to influence hand selection (Schweighofer et al. 2015). The relative effort associated with moving either arm changes continuously under whole body motion, which we have hypothesized might alter hand choice during passive body motion (Bakker et al. 2017). Bakker et al. (2017) found that a model that computes future movement effort based on a constant whole-body acceleration from the moment of target presentation, best describes the observed choice biases. Behaviorally we confirm previous results, but the phase shift between behavior and corticospinal excitability in the current study suggests that the exact moment at which a snapshot of the body acceleration is taken might be later than the moment of target presentation.

From previous work we know that reaching under whole body linear acceleration requires adaptation of an internal model (Sarwary et al. 2013), but we were unable to show signs of learning in choice behavior in the current paradigm (Bakker et al. 2017). This raises the question whether the modulation of the MEP with sled phase is developing over the course of the experiment or that this coupling is the result of a more direct modulation of body-motion related signals on the corticospinal tract. We tried to post-hoc examine the possibility of a development of the MEP amplitude due to learning by comparing the MEPs of the first half of the experiment to the second half, whereby MEP amplitude was computed for each half separately according to the methods applied to the full dataset. However, the small number of trials did not provide us with enough power to prove or disprove a change in MEP over the course of the experiment. Future research, with more TMS trials, might examine if corticospinal excitability slowly adapts as a function of the passive body motion.

The observed modulation in corticospinal excitability may find its origin in cortical areas related to hand selection and movement preparation, but also in the spinal part of the circuit (Bestmann and Krakauer 2015). Possibly postural responses anticipating the passive full-body motion modulated the MEP amplitude (Kazennikov et al. 2005). To minimize co-activation of antagonistic muscle pairs (Selen et al. 2005), our set-up was designed to enable a relaxed arm and body posture throughout the experiment. If TLAT was unexpectedly more active than during resting state, this trial was excluded. Also, if there would have been co-activation, one may expect that this would result in an overall increase of muscle tension, rather than the sinusoidal pattern observed here. Therefore, we believe that the observed modulation of corticospinal excitability with body motion is not related to increased tension in antagonistic muscles, but rather to the motion itself.

Alternatively, more global effects could have influenced corticospinal activity. For example, it has been reported that attentional focus (external versus internal) modulates MEPs evoked by motor cortex stimulation (Kuhn et al. 2018). Concurrent leg muscle activation results in a prolonged attenuation of EMG activity (i.e. cortical silent period) after TMS pulses targeting finger muscle abduction, while the MEP amplitude itself remained unaffected (Sohn et al. 2005). Bestmann et al. (Bestmann et al. 2008) demonstrated that uncertainty and surprise influence MEPs in a delayed-response task. Although our TMS pulses were applied over M1, the induced electric field could have resulted in stimulation of cortico-spinal, intra-cortical and trans-cortical neurons, with activation spreading throughout the cerebral cortex possibly increasing neural excitability (Bestmann and Krakauer 2015; Casula et al. 2018). We controlled for these effects by means of full body fixation, no target being present in the TMS trials and unpredictable stimuli presentation times.

We manipulated the state of the body with sinusoidal full-body motion. Since position, velocity and acceleration are inherently related for sinusoids, and the motion may be predictable, it is difficult to conclude what information participants used. Congruent with previous findings for eye and hand selection, peak preferences align with acceleration information (Bakker et al. 2017, 2019; Rincon-Gonzalez et al. 2016). However, corticospinal excitability peaks around phases of maximum and minimum velocity. Future work could use multiple superimposed sinusoidal sled motions, whereby position, velocity and acceleration are decoupled, to test what information drives behavior and corticospinal excitability.

To conclude, we show that both choice behavior and corticospinal excitability modulate as a function of passive full body motion. This modulation may be driven by biomechanical costs predicted based on vestibular information, suggesting that body motion information biases hand selection processes even before a target is presented.

## Data Availability Statement

Upon publication, all data and code are available from the Donders Institute for Brain, Cognition and Behaviour repository at: https://doi.org/10.34973/pm0w-wj61

## Authors Contributions

LOW, BSR, DJLGS, LPJS and WPM conception and design of research; LOW, SCW, BSR and LPJS performed experiments; LOW and LPJS analyzed data; LOW, LPJS and WPM interpreted results of experiments; LOW prepared figures; LOW drafted manuscript; LOW, SCW, BSR, DJLGS, LPJS and WPM edited and revised manuscript; LOW, SCW, BSR, DJLGS, LPJS and WPM approved final version of manuscript.

## References

Angelaki DE, Cullen KE. Vestibular System: The Many Facets of a Multimodal Sense. Annual Review of Neuroscience 31: 125–150, 2008.

Bakker RS, Selen LPJ, Medendorp WP. Reference frames in the decisions of hand choice. J Neurophysiol 119: 1809–1817, 2018.

Bakker RS, Selen LPJ, Medendorp WP. Transformation of vestibular signals for the decisions of hand choice during whole body motion. J Neurophysiol 121: 2392–2400, 2019.

Bakker RS, Weijer RHA, van Beers RJ, Selen LPJ, Medendorp WP. Decisions in motion: passive body acceleration modulates hand choice. Journal of Neurophysiology 117: 2250–2261, 2017.

Berens P. CircStat: A MATLAB Toolbox for Circular Statistics. Journal of Statistical Software 31, 2009.

Bernier P-M, Cieslak M, Grafton ST. Effector selection precedes reach planning in the dorsal parietofrontal cortex. Journal of Neurophysiology 108: 57–68, 2012.

Bestmann S, Harrison LM, Blankenburg F, Mars RB, Haggard P, Friston KJ, Rothwell JC. Influence of uncertainty and surprise on human corticospinal excitability during preparation for action. Curr Biol 18: 775–780, 2008.

Bestmann S, Krakauer JW. The uses and interpretations of the motor-evoked potential for understanding behaviour. Exp Brain Res 233: 679–689, 2015.

Bresciani J-P, Gauthier GM, Vercher J-L, Blouin J. On the nature of the vestibular control of arm-reaching movements during whole-body rotations. Experimental Brain Research 164: 431–441, 2005.

Casula EP, Rocchi L, Hannah R, Rothwell JC. Effects of pulse width, waveform and current direction in the cortex: A combined cTMS-EEG study. Brain Stimulation 11: 1063–1070, 2018.

Cisek P. Cortical mechanisms of action selection: the affordance competition hypothesis. Philos Trans R Soc Lond, B, Biol Sci 362: 1585–1599, 2007.

Cisek P, Kalaska JF. Neural correlates of reaching decisions in dorsal premotor cortex: specification of multiple direction choices and final selection of action. Neuron 45: 801–814, 2005.

Cisek P, Kalaska JF. Neural mechanisms for interacting with a world full of action choices. Annu Rev Neurosci 33: 269–298, 2010.

Cos I, Belanger N, Cisek P. The influence of predicted arm biomechanics on decision making. Journal of Neurophysiology 105: 3022–3033, 2011.

Cos I, Duque J, Cisek P. Rapid prediction of biomechanical costs during action decisions. J Neurophysiol 112: 1256–1266, 2014.

Dekleva BM, Kording KP, Miller LE. Single reach plans in dorsal premotor cortex during a two-target task. Nat Commun 9: 3556, 2018.

Dekleva BM, Ramkumar P, Wanda PA, Kording KP, Miller LE. Uncertainty leads to persistent effects on reach representations in dorsal premotor cortex. eLife 5: e14316, 2016.

Derosiere G, Thura D, Cisek P, Duque J. Motor cortex disruption delays motor processes but not deliberation about action choices. Journal of Neurophysiology 122: 1566–1577, 2019.

Duque J, Ivry RB. Role of corticospinal suppression during motor preparation. Cereb Cortex 19: 2013–2024, 2009.

Duque J, Lew D, Mazzocchio R, Olivier E, Ivry RB. Evidence for two concurrent inhibitory mechanisms during response preparation. J Neurosci 30: 3793–3802, 2010.

Kass RE, Raftery AE. Bayes Factors. Journal of the American Statistical Association 90: 773–795, 1995.

Kazennikov O, Solopova I, Talis V, Grishin A, Ioffe M. TMS-responses during anticipatory postural adjustment in bimanual unloading in humans. Neuroscience Letters 383: 246–250, 2005.

Keyser J, Medendorp WP, Selen LPJ. Task-dependent vestibular feedback responses in reaching. Journal of Neurophysiology 118: 84–92, 2017.

Klein-Flügge MC, Bestmann S. Time-dependent changes in human corticospinal excitability reveal value-based competition for action during decision processing. J Neurosci 32: 8373–8382, 2012.

Klein-Flügge MC, Nobbs D, Pitcher JB, Bestmann S. Variability of human corticospinal excitability tracks the state of action preparation. J Neurosci 33: 5564–5572, 2013.

Kontsevich LL, Tyler CW. Bayesian adaptive estimation of psychometric slope and threshold. Vision Research 39: 2729–2737, 1999.

Kuhn Y-A, Keller M, Lauber B, Taube W. Surround inhibition can instantly be modulated by changing the attentional focus. Sci Rep 8: 1085, 2018.

Mamolo CM, Roy EA, Bryden PJ, Rohr LE. The effects of skill demands and object position on the distribution of preferred hand reaches. Brain Cogn 55: 349–351, 2004.

Oliveira FTP, Diedrichsen J, Verstynen T, Duque J, Ivry RB. Transcranial magnetic stimulation of posterior parietal cortex affects decisions of hand choice. Proc Natl Acad Sci USA 107: 17751–17756, 2010.

Oostwoud Wijdenes L, van Beers RJ, Medendorp WP. Vestibular modulation of visuomotor feedback gains in reaching. Journal of Neurophysiology 122: 947–957, 2019.

Prins N. The psi-marginal adaptive method: How to give nuisance parameters the attention they deserve (no more, no less). Journal of Vision 13: 3–3, 2013.

Przybyla A, Coelho CJ, Akpinar S, Kirazci S, Sainburg RL. Sensorimotor performance asymmetries predict hand selection. Neuroscience 228: 349–360, 2013.

Ratcliff R, McKoon G. The Diffusion Decision Model: Theory and Data for Two-Choice Decision Tasks. Neural Computation 20: 873–922, 2008.

Reichenbach A, Bresciani J-P, Bülthoff HH, Thielscher A. Reaching with the sixth sense: Vestibular contributions to voluntary motor control in the human right parietal cortex. NeuroImage 124: 869–875, 2016.

Rincon-Gonzalez L, Selen LPJ, Halfwerk K, Koppen M, Corneil BD, Medendorp WP. Decisions in motion: vestibular contributions to saccadic target selection. J Neurophysiol 116: 977–985, 2016.

Rossi S, Hallett M, Rossini PM, Pascual-Leone A. Safety, ethical considerations, and application guidelines for the use of transcranial magnetic stimulation in clinical practice and research. Clinical Neurophysiology 120: 2008–2039, 2009.

Sarwary AME, Selen LPJ, Medendorp WP. Vestibular benefits to task savings in motor adaptation. J Neurophysiol 110: 1269–1277, 2013.

Schutter DJLG, van Honk J. A Standardized Motor Threshold Estimation Procedure for Transcranial Magnetic Stimulation Research: The Journal of ECT 22: 176–178, 2006.

Schweighofer N, Xiao Y, Kim S, Yoshioka T, Gordon J, Osu R. Effort, success, and nonuse determine arm choice. J Neurophysiol 114: 551–559, 2015.

Selen LPJ, Beek PJ, Dieën JH van. Can co-activation reduce kinematic variability? A simulation study. Biol Cybern 93: 373–381, 2005.

Sohn YH, Kang SY, Hallett M. Corticospinal disinhibition during dual action. Exp Brain Res 162: 95–99, 2005.

Stoloff RH, Taylor JA, Xu J, Ridderikhoff A, Ivry RB. Effect of reinforcement history on hand choice in an unconstrained reaching task. Front Neurosci 5: 41, 2011.

Thura D, Cisek P. Deliberation and commitment in the premotor and primary motor cortex during dynamic decision making. Neuron 81: 1401–1416, 2014.

Wichmann FA, Hill NJ. The psychometric function: I. Fitting, sampling, and goodness of fit. Perception & Psychophysics 63: 1293–1313, 2001.

